# Stage-specific disruption of X chromosome expression during spermatogenesis in sterile house mouse hybrids

**DOI:** 10.1101/2021.11.12.468424

**Authors:** Erica L. Larson, Emily E. K. Kopania, Kelsie E. Hunnicutt, Dan Vanderpool, Sara Keeble, Jeffrey M. Good

## Abstract

Hybrid sterility is a complex phenotype that can result from the breakdown of spermatogenesis at multiple developmental stages. Here, we disentangle two proposed hybrid male sterility mechanisms in the house mice, *Mus musculus domesticus* and *M. m. musculus*, by comparing patterns of gene expression in sterile F1 hybrids from a reciprocal cross. We found that hybrid males from both cross directions showed disrupted X chromosome expression during prophase of meiosis I consistent with a loss of Meiotic Sex Chromosome Inactivation (MSCI) and *Prdm9*-associated sterility, but that the degree of disruption was greater in mice with an *M. m. musculus* X chromosome consistent with previous studies. During postmeiotic development, gene expression on the X chromosome was only disrupted in one cross direction, suggesting that misexpression at this later stage was genotype-specific and not a simple downstream consequence of MSCI disruption which was observed in both reciprocal crosses. Instead, disrupted postmeiotic expression may depend on the magnitude of earlier disrupted MSCI, or the disruption of particular X-linked genes or gene networks. Alternatively, only hybrids with a potential deficit of *Sly* copies, a Y-linked ampliconic gene family, showed overexpression in postmeiotic cells, consistent with a previously proposed model of antagonistic coevolution between the X and Y-linked ampliconic genes contributing to disrupted expression late in spermatogenesis. The relative contributions of these two regulatory mechanisms and their impact on sterility phenotypes awaits further study. Our results further support the hypothesis that X-linked hybrid sterility in house mice has a variable genetic basis, and that genotype-specific disruption of gene regulation contributes to overexpression of the X chromosome at different stages of development. Overall, these findings underscore the critical role of epigenetic regulation of the X chromosome during spermatogenesis and suggest that these processes are prone to disruption in hybrids.

## INTRODUCTION

Hybrid sterility can result from the breakdown of gametogenesis at several developmental stages, from early divisions of mitotic cells, meiosis, to the differentiation of postmeiotic cells into mature gametes. After gamete production, hybrid fertility can also be reduced through mechanisms that impede fertilization, such as a failure of hybrid sperm to transfer or fertilize. In hybrid males, sterility is typically measured by quantitative traits such as testes weight and histology; sperm counts, motility, and morphology; and the ability to sire offspring. Often, these traits are correlated (White *et al.* 2011; Turner and Harr 2014; Larson *et al.* 2018b) and are evaluated as though they were a single phenotype, but that does not mean sterility arises from a single mechanism or genetic basis (Reed and Markow 2004; Campbell and Nachman 2014). To tease apart different mechanisms of hybrid sterility requires a developmental framework, where breakdown at different stages of spermatogenesis can be evaluated to understand, as a whole, the evolution of hybrid sterility (Larson *et al.* 2018a; Cutter and Bundus 2020).

Hybrid sterility is often a composite phenotype because it typically has a complex genetic basis that involves the negative epistatic interactions of multiple alleles, known as Dobzhansky-Muller Incompatibilities or DMIs (Dobzhansky 1937; Muller 1942, see also Bateson 1909). Incompatible alleles can be polymorphic or have modifiers that affect their expression (Cutter 2012), so that the extent of reproductive isolation varies among individuals within or between populations (Good *et al.* 2008b; Sweigart and Flagel 2015; Case *et al.* 2016; Mandeville *et al.* 2017; Bracewell *et al.* 2017; Zuellig and Sweigart 2018). DMIs can also evolve early in the divergence process (Coughlan and Matute 2020) and are expected to accumulate over time so that many different epistatic combinations of alleles may contribute to hybrid breakdown (Moyle and Nakazato 2010; Wang *et al.* 2013). Gene flow between populations can also lead to recombination of incompatible alleles (Bank *et al.* 2012; Lindtke and Buerkle 2015), which can further complicate patterns of population-level variation in DMIs (Larson *et al.* 2018b; Meiklejohn *et al.* 2018). For all these reasons, careful laboratory dissection of sterility phenotypes remains a critical component of understanding the genetic basis of speciation.

Between subspecies of house mice, *Mus musculus domesticus* and *M. m. musculus* (hereafter *domesticus* and *musculus*), the evolution of hybrid sterility appears to be due to a combination of several genetic factors. These subspecies diverged ~350-500 mya (Geraldes *et al.* 2011; Duvaux *et al.* 2011; Phifer-Rixey *et al.* 2020) and have come into secondary contact in a long hybrid zone in central Europe (Macholán *et al.* 2012; Phifer-Rixey and Nachman 2015). Female hybrids are generally more fertile than males (but see Suzuki and Nachman 2015) and hybrid male fertility varies considerably in the hybrid zone (Turner *et al.* 2012). In crosses between *domesticus* females and *musculus* males, hybrid male sterility depends on which individual genotypes are sampled, while crosses between *musculus* females and *domesticus* males typically produce sterile hybrid males (Vanlerberghe *et al.* 1986; Alibert *et al.* 1997; Britton-Davidian *et al.* 2005; Vyskočilová *et al.* 2005; Good *et al.* 2008b; Turner *et al.* 2012).

There are many different autosomal regions that have been associated with hybrid sterility in house mice (*e.g.,* Oka *et al.* 2007; Good *et al.* 2008a; White *et al.* 2011; Turner *et al.* 2014; Turner and Harr 2014; Larson *et al.* 2018b; Schwahn *et al.* 2018; Morgan *et al.* 2020; Widmayer *et al.* 2020), but the primary genetic determinant of sterility in F1 hybrid males involves the rapid evolution of PRDM9 binding sites, the autosomal encoded protein that directs the location of recombination in mammals (Mihola *et al.* 2009; Mukaj *et al.* 2020). In F1 mouse hybrids, PRDM9 binds preferentially to ancestral binding sites, leading to the asymmetric formation of double strand breaks and autosomal asynapsis (Davies *et al.* 2016; Gregorova *et al.* 2018). When the number of asynapsed chromosomes in a cell reaches a threshold, it can trigger cell death and in the most severe cases, complete meiotic arrest (Bhattacharyya *et al.* 2013). *Prdm9*-associated sterility is polymorphic, with alternative ‘fertile’ *Prdm9* alleles (Flachs *et al.* 2012; Mukaj *et al.* 2020) and is further modulated by epistatic interactions with a locus on the *musculus* X chromosome (*Hstx2,* Storchová *et al.* 2004; Bhattacharyya *et al.* 2014; Lustyk *et al.* 2019). A characteristic signal of *Prdm9-*associated sterility is the overexpression of the X chromosome during early meiosis I (Good *et al.* 2010; Bhattacharyya *et al.* 2013; Campbell *et al.* 2013; Turner *et al.* 2014; Larson *et al.* 2017), a developmental stage where the X chromosome would normally be transcriptionally inactive known as Meiotic Sex Chromosome Inactivation (MSCI, Turner 2015). Whether disrupted MSCI is a byproduct, or an integral part of *Prdm9*-associated sterility is still unclear (Forejt *et al.* 2021), but it is a distinct regulatory phenotype of hybrid sterility at this developmental stage.

Hybrid male sterility in house mice may also be influenced by interactions among three ampliconic sex-linked gene families expressed in postmeiotic cells, *Slx* and *Slxl1* (X chromosome) and *Sly* (Y chromosome, Ellis *et al.* 2011; Cocquet *et al.* 2012). SLY plays a central role in repressing the transcription of sex-linked genes, known as postmeiotic sex chromosome repression (PSCR), while SLX/SLXL1 counteract the repression of SLY by competing for binding access to SSTY1 at the promoter of thousands of postmeiotic genes (Moretti *et al.* 2020). *SLY* and *SLX/SLXL1* appear to compete through a copy-number arms race, with higher relative gene copies of *Sly* leading to the repression of other multicopy genes. Gene knockdowns of *Sly* (*i.e.*, *Sly*-deficient) result in the overexpression of the X chromosome and female-biased litters (Cocquet *et al.* 2009; Kruger *et al.* 2019), while knockdowns of *Slx*/*Slxl1* (*i.e.*, *Slx*-deficient) result in a slight underexpression of the X chromosome and male-biased litters (Cocquet *et al.* 2010, 2012; Kruger *et al.* 2019). These genes have undergone a massive co-amplification across different mouse lineages, leading to different copy numbers in *domesticus* and *musculus* (Ellis *et al.* 2011; Morgan and Pardo-Manuel de Villena 2017). As a result, F1 hybrids between *musculus* females and *domesticus* males potentially have a deficit of *Sly* gene copies, while hybrids from the reciprocal cross have a deficit of *Slx/Slxl1* gene copies (Good 2012). We previously demonstrated that the X chromosome in postmeiotic cells is overexpressed in *Sly*-deficient hybrids, consistent with *Sly*/*Slx*-associated sterility (Larson *et al.* 2017). We also observed overexpression in *Sly*-deficient hybrids of an ampliconic autosomal gene family, *α-takusan*, that is regulated by *SLY* (Moretti *et al.* 2017) and a slight underexpression of the X chromosome in *Slx*-deficient hybrids, consistent with *Sly* repression (Kruger *et al.* 2019). These results support a model of postmeiotic disruption of X chromosome expression and *Sly*/*Slx*-associated sterility.

Incompatibilities at each of these stages may produce similar sterility phenotypes, such as low testes weight and abnormal sperm morphology, making it difficult to tease apart their contribution to overall hybrid sterility and the maintenance of the house mouse hybrid zone. The disrupted expression of the X chromosome at different developmental stages suggests that hybrid sterility in these mice is a composite of multiple regulatory mechanisms (Larson *et al.* 2017). However, because both *Prdm9* and *Sly*/*Slx* associated sterility are often asymmetric and depend on interactions with the *M. m. musculus* X chromosome it is possible that postmeiotic disruption of the X chromosome observed in some crosses is simply a downstream effect of disrupted MSCI and a cascade of disrupted X chromosome expression. In this study, we used an independent cross to help disentangle the effects of regulatory disruption at different developmental stages of spermatogenesis. We used strains of mice that produce subfertile hybrid males in both cross directions, but only offspring from *musculus* females and *domesticus* males have a *Sly* deficit. We found that both reciprocal hybrids showed disrupted MSCI, consistent with *Prdm9*-associated sterility. However, only the hybrids that had the greater disruption of MSCI and are *Sly*-deficient showed disrupted postmeiotic X chromosome expression, suggesting that postmeiotic disruption is genotype-specific. Collectively, these results further underscore the considerable genotypic and phenotypic (regulatory and reproductive) variably underlying F1 hybrid sterility between these closely-related mouse lineages.

## MATERIALS & METHODS

### Crosses and reproductive phenotypes

We used four inbred strains of wild-derived mice from two subspecies of *domesticus* (WSB/EiJ and LEWES/EiJ) and *musculus* (PWK/PhJ and CZECHII/EiJ). First, we generated intraspecific F1s between strains of *domesticus* (WSB females × LEWES males) and *musculus* (CZECHII females × PWK males). These mice served as parental controls for each species, but without the negative effects of inbreeding on male fertility. Second, we generated intersubspecific F1 hybrids in reciprocal crosses between one strain of each subspecies (CZECHII females x WSB males and WSB females x CZECHII males, *hereafter*: ♀*mus*^CZII^ × ♂*dom*^WSB^ and ♀*dom*^WSB^ × ♂*mus*^CZII^; throughout the manuscript we will indicate all crosses as female parent × male parent). We chose crosses involving CZECHII mice because F1 hybrid males from these crosses are subfertile in both directions of the cross (Good *et al.* 2008b; Larson *et al.* 2018b). This provided a direct contrast to other studies using strains that produce subfertile F1 hybrid males in only one cross direction, (*i.e.* PWK females x LEWES males, Good *et al.* 2010; Campbell *et al.* 2013; Mack *et al.* 2016; Larson *et al.* 2017), which allowed us to begin to isolate the effects of disrupted MSCI and imbalanced copy numbers of *Slx* and *Sly* on regulatory phenotypes. Experimental mice used in this study were obtained from breeding colonies established from mice purchased from The Jackson Laboratory (Bar Harbor, ME) in 2010 and were maintained at the University of Montana Department of Laboratory Animal Resources (IACUC protocol 002-13). One *domesticus* mouse had a sire from replacement stock of LEWES/EiJ ordered in 2013. The stock origin for each mouse is indicated in **Table S1**.

We weaned experimental mice at ~21 days after birth and housed them in same sex sibling groups until males were individually isolated at 45 days. We euthanized males between 61 and 89 days old using CO_2_ followed by cervical dislocation. Immediately after euthanasia we quantified male reproductive traits following previously described protocols (Good *et al.* 2008b, 2008a). We weighed paired testes and seminal vesicles relative to body weight and isolated sperm by dicing the caudal epididymides in 1 mL of Dulbecco’s PBS (Sigma) followed by a 10 min incubation at 37°C. We estimated the proportion of motile sperm and total sperm numbers using 5 μL sperm suspensions (regular and heat-shocked, respectively) viewed in a Makler counting chamber on a light microscope over a fixed area and observation time. We fixed and stained 25 μL sperm suspensions and later counted 100 intact sperm to visually classify morphology. All samples were counted by a single individual (E.L.L.) while blind to genotype. We classified sperm as (1) normal with a long apical hook, (2) slightly abnormal with a shortened hook, (3) abnormal with a short hook and rounded shape, and (4) severely abnormal with an amorphous shape. We summarized sperm morphology using a weighted index that ranged from high (3) to low (0) following Oka *et al.* (2004) and Good *et al.* (2008a).

### RNA sequencing of spermatogenesis stages

Testes are composed of at least eleven major cell types, with cell-specific patterns of gene expression (Margolin *et al.* 2014; Green *et al.* 2018). Whole testes expression patterns can be confounded by differences in cell composition between species, or between sterile and fertile hybrids (Good *et al.* 2010; Hunnicutt *et al.* 2021). To overcome these challenges, we used fluorescence activated cell sorting (FACS) to isolate highly enriched cell populations for three developmental stages of spermatogenesis: early prophase of meiosis I prior to MSCI (leptotene/zygotene cells), meiosis I after MSCI (diplotene cells) and postmeiotic development prior to spermiogenesis (round spermatids). Our complete FACS protocol, modified from Getun *et al.* (2011), is available on Github (https://github.com/goodest-goodlab/good-protocols/tree/main/protocols/FACS). We decapsulated the testes and disassociated them in a mixture of 1 mg/mL collagenase (Worthington Biochemical), GBSS (Sigma) and 1 mg/mL trypsin (Worthington Biochemical). We inactivated the trypsin with 0.16 mg/mL fetal calf serum (Sigma) and stained the cells with 0.36 mg/mL of Hoechst 33343 (Invitrogen) and 0.002 mg/mL propidium iodide. At each step, we incubated solutions in a mini shaker at 120 rpm at 33°C for 15 min and added 0.004 mg/mL DNase to eliminate clumps. We filtered disassociated cells twice using a 40 μm strainer and sorted cells on a FACSAria IIu cell sorter (BD Biosciences) at the UM Center for Environmental Health Sciences Fluorescence Cytometry Core. FACS isolates cells based on size, granularity, and fluorescence (traits that change across different stages of spermatogenesis). We collected enriched cell populations in 15 μL beta mercaptoethanol (Sigma) per mL of RLT lysis buffer (Qiagen) and extracted RNA from each cell type using a Qiagen RNeasy kit. We quantified our samples on a Bioanalzyer 2000 (Agilent) and prepared samples with RNA integrity (RIN) above 8 for sequencing using an Illumina Truseq Sample Prep Kit v2 in a design that avoided batch effects between cell populations and genotypes. We extracted RNA from a total of 21 mice, using the highest quality enriched cell populations to generate RNAseq libraries for three individuals per cell type, three cell types and four crosses (*domesticus*, *musculus* and their reciprocal F1 hybrids, *n* = 36 RNAseq libraries).

We sequenced each library on an Illumina HiSeq 2500 (SE, 100 bp) at the University of Oregon Genomics and Cell Characterization Core Facility and on an Illumina HiSeq 2000 (PE, 100 bp) and a NextSeq 500 (SE, 100 bp) at the University of Southern California Epigenome Center. While all of the RNAseq libraries in this study were prepared simultaneously, we previously published a subset of these data, the *domesticus* and *musculus* parent samples, as part of a study on the rate of molecular evolution in spermatogenesis (Larson *et al.* 2016). Here we focus on comparisons between reciprocal F1 hybrids (unpublished data) and their parents, to disentangle the effects of different developmental stages on regulatory disruption in hybrids.

### Read mapping and differential expression analyses

We trimmed reads using TRIMMOMATIC v0.32 (Bolger *et al.* 2014) and mapped reads using TOPHAT v2.0.10 (Kim *et al.* 2013) to strain-specific pseudo-references for *domesticus* (WSB/EiJ) and *musculus* (PWK/PhJ) (Huang *et al.* 2014). These pseudo-references incorporate all known SNPs, indels and structural variants for these strains relative to the Genome Reference Consortium mouse build 38 (GRCm38), thereby minimizing mapping bias to the mouse reference genome, which is predominately *domesticus* (Yang *et al.* 2011). We used LAPELS v1.0.5 to translate our reads back into the GRCm38 coordinates and SUSPENDERS v0.2.4 to merge our alignments (Huang *et al.* 2014). We counted the number of reads that mapped to protein-coding genes (Ensembl release 78) using FEATURECOUNTS v1.4.4 (Liao *et al.* 2014). We counted reads that were 1) uniquely mapped to a single protein-coding gene and 2) mapped to multiple protein-coding genes. These two approaches were qualitatively the same, but by including multi-mapped reads we could account for the expression of multicopy gene families that are enriched on the mouse X chromosome, and in all cases we report these results.

We analyzed gene expression using Bioconductor v3.0 package edgeR v3.30.3 (Robinson *et al.* 2010) in R v4.0.1 (R Core Team 2020). We normalized our data using the scaling factor method and restricted our analysis to genes with a minimum expression of FPKM > 1 in at least three samples. For all analyses, we tested alternative normalization methods (*e.g.,* weighted trimmed mean of M-values) and found qualitatively similar results. We fit our data with a negative binomial generalized linear model with Cox-Reid tagwise dispersion estimates (McCarthy *et al.* 2012). Our model included cross and cell type as a single factor and our design matrix contrasted different crosses for each cell type. To evaluate differential expression, we used likelihood ratio tests, dropping one coefficient from the design matrix and comparing that to the full model. For each contrast, we restricted our differentially expressed (DE) genes to genes that are expressed in the focal cell type (FPKM > 1 in 3/6 samples) and in all cases used a p-value adjusted for a false discovery rates (FDR) of 5% (Benjamini and Hochberg 1995). For all our RNAseq analysis, we focused on contrasts between each hybrid and their parental X chromosome, to account for potential mapping biases on the hemizygous X (♀*mus*^CZII^ × ♂*dom*^WSB^ vs. *musculus*; ♀*dom*^WSB^ × ♂*mus*^CZII^ vs. *domesticus*) and contrasts between the two F1 hybrids (♀*mus*^CZII^ × ♂*dom*^WSB^ vs. ♀*dom*^WSB^ × ♂*mus*^CZII^).

We used a sliding gene window to test for local enrichment of autosomal genes that were overexpressed in round spermatids of ♀*mus*^CZII^ × ♂*dom*^WSB^ hybrids compared to ♀*dom*^WSB^ × ♂*mus*^CZI^ hybrids. We counted the proportion of genes that were up (+logFC) or down (−logFC) regulated within a given window and identified windows that fell outside of the 99^th^ quantile modeled with a Poisson distribution. We tested a range of window sizes (50-400 genes/window) and found qualitatively similar results, so we used 250 genes/window. This method has been previously used to identify overexpression of an ampliconic autosomal gene family, *α-takusan* in sterile *musculus* x *domestics* hybrids (Larson et al. 2017).

### Sequencing of *Prdm9* alleles

We characterized the *Prdm9* Exon12 allele for each strain used in our study. For each strain, we extracted DNA from liver tissue of a single mouse using a Nucleospin Tissue Kit (Macherey-Nagel) and quantified the DNA with a QuantiFluor dsDNA System (Promega) on a Synergy HTX Multi-Mode Microplate Reader (Agilent). We amplified *Prdm9* Exon12 using the primers Exon12-L1 and Exon12-R (Mukaj *et al.* 2020), GoTaq Polymerase (Promega) and the following protocol: an initial denaturation at 95°C for 2 min, followed by 41 cycles of 95°C for 30 s, 56°C for 30 s, and 72°C for 1 min, with a final extension step of 72°C for 5 min. We purified and sequenced amplicons at Genewiz (New Jersey, USA), using their hairpin sequencing. We manually cleaned and translated sequences in Geneious 9.1.8 (Biomatters) and aligned sequences using MAFFT v7.453 (Katoh and Standley 2013). We identified C-terminal zinc finger domains by searching sequences with hmmsearch for the Zf-C2H2 HMM profile (PF00096.27) from the Pfam database (HMMER v3.3.2; Mistry *et al.* 2021). We excluded the first nonvariant zinc finger domain then compared the −1, 3, and 6 positions within each domain (as in Oliver *et al.* 2009) to previously published *musculus* and *domesticus* alleles (Mukaj *et al.* 2020).

### Gene Copy Number Estimates

To estimate *Slx* and *Sly* gene copy numbers, we generated whole genome sequence data from a single mouse of each strain used in our study (*domesticus* WSB/EiJ, LEWES/EiJ and *musculus* PWK/PhJ and CZECHII/EiJ). For each sample, we prepared and sequenced libraries twice to increase unique read coverage. We extracted DNA from liver tissue using a Qiagen DNeasy kit and sent samples to Novogene for library preparation and sequencing on an Illumina NovaSeq 6000 (PE, 150bp). We trimmed reads with TRIMMOMATIC v0.39 (Bolger *et al.* 2014), mapped our reads to the GRCm38 using BWA-MEM v0.7.17 (Li and Durbin 2009), and fixed mates and marked duplicates with Picard v2.18.29 (Broad Institute 2019). We merged the data from each sequencing effort resulting in 10-15X average genome-wide coverage.

To identify paralogs of ampliconic gene families, we extracted *Slx*, *Slxl1*, and *Sly* gene sequences from the mouse reference GRCm38 using Ensembl release 102 (Yates *et al.* 2019). We performed Ensembl BLAT searches with these sequences against the GRCm38 mouse reference, allowing up to 1000 hits. We then extracted all BLAT hits with greater than or equal to 97% sequence identity and an e-value of 0.0 and considered these filtered BLAT hits to be gene family paralogs for downstream copy number estimation.

We estimated copy number using two methods based on relative coverage. First, we followed a similar approach as Morgan and Pardo-Manuel de Villena (2017) and used Mosdepth (Pedersen and Quinlan 2018) to estimate coverage in paralog regions and the average coverage across the whole genome. We estimated copy number by summing coverage across paralog regions and dividing this sum by half the genome-wide average coverage. We halved the average coverage because most of the mouse genome is diploid, while the sex chromosomes in males are haploid. We also used the approach implemented in AmpliCoNE (Vegesna *et al.* 2019), which estimates copy number from relative coverage using only regions that are considered informative based on repeat masking and mappability, while also controlling for GC content. AmpliCoNE was developed for estimating gene copy numbers on the human Y chromosome, so we made some modifications to account for the less complete assembly and annotation of the mouse sex chromosomes. Specifically, instead of relying on informative sites to differentiate copy numbers, we extracted all kmers of length 101bp from the *Slx*, *Slxl1*, and *Sly* gene sequences and mapped these back to the mouse reference genome using Bowtie2, allowing up to 500 multiple mapping hits. For each gene, we identified the most frequent number of times (*m*) kmers mapped to the mouse genome and kept only kmers that mapped *m* times. We identified all locations where these kmers mapped with 2 or fewer mismatches and used these kmer start locations as the “informative sites” metric for AmpliCoNE.

### Data Availability

The data reported in this paper are available through the National Center for Biotechnology Information Sequence Read Archive under accession numbers PRJNA296926 (*domesticus* and *musculus* RNAseq data), PRJNA352861 (F1 hybrid RNAseq data), PRJNA732719 (lab strain whole genome sequence data). *Prdm9* sequences were deposited in Genbank under accession numbers MZ733983-MZ733986. Male reproductive phenotype data are available in **Table S1**.

## RESULTS

### Hybrid males from both cross directions were subfertile

We found F1 hybrid males from crosses between *domesticus* (WSB) and *musculus* (CZECHII) were subfertile in both cross directions, but that ♀*mus*^CZII^ × ♂*dom*^WSB^ hybrids had more severe abnormal sperm morphology. Overall, ♀*dom*^WSB^ × ♂*mus*^CZII^ hybrids had lower fertility than both *domesticus* and *musculus*, with significantly smaller testes, lower sperm counts, and more abnormal sperm morphology, while ♀*mus*^CZII^ × ♂*dom*^WSB^ hybrids had smaller testes and lower sperm counts, but after correcting for multiple tests these values were only significant in comparisons with *domesticus* (**Table 1**). The ♀*mus*^CZII^ × ♂*dom*^WSB^ hybrids did have the most severely abnormal sperm morphology consistent with previous studies (Good *et al.* 2008b; Larson *et al.* 2018b). There were no significant differences in the relative seminal vesicle weight or the proportion of motile sperm across any comparisons.

**Table 1.** Reproductive phenotypes of male mice used in this study. The table summarizes the sample sizes for each cross (N) and the median (± standard error) trait value for five reproductive phenotypes. Arrows indicate whether the hybrids had significantly lower reproductive values relative to *domesticus* (closed arrows) or *musculus* (open arrows). Values in bold indicate traits that were significantly different between the two F1 hybrids. Testes and seminal vesicle weights are reported relative to body size. The sperm morphology index ranged from 3 (high quality sperm) to 0 (severally abnormal sperm). Significance was estimated using a Wilcoxon test with p-values FDR corrected for multiple comparisons.

| Cross | N | Relative testis weight (mg/g) | Relative seminal vesicle weight (mg/g) | Proportion motile sperm | Sperm count (1x10<sup>6</sup>) | Sperm head morphology index |
| --- | --- | --- | --- | --- | --- | --- |
| *domesticus* | 6 | 11.30 ± 0.34 | 5.18 ± 0.29 | 0.82 ± 0.04 | 14.8 ± 1.80 | 2.99 ± 0.01 |
| *musculus* | 5 | 9.53 ± 0.63 | 5.58 ± 1.30 | 0.87 ± 0.06 | 17.8 ± 2.50 | 3.00 ± 0.03 |
| ♀ *dom*<sup>WSB</sup> × ♂ *mus*<sup>CZII</sup> | 6 | ▼▽6.28 ± 0.28 | 5.68 ± 0.54 | 0.83 ± 0.06 | ▼▽4.2 ± 0.72 | ▼▽1.29 ± 0.07 |
| ♀ *mus*<sup>CZII</sup> × ♂ *dom*<sup>WSB</sup> | 4 | ▼6.46 ± 0.35 | 5.44 ± 0.27 | 0.65 ± 0.12 | ▼5.8 ± 2.70 | ▼▽0.66 ± 0.12 |

### Cell-specific gene expression

For each cross, we generated between 14.7 and 26.8 million mapped fragments (paired or unpaired reads) per cell type (738 million total, mean = 20.5 million). After filtering we retained 14,209 expressed protein-coding genes. Gene expression profiles clustered by cell type (**Fig S1A**) and within each cell type, samples clustered by parental species with F1 hybrids intermediate to the two parents (**Fig S1B–D**). Overall, the strong clustering by cell type and cross, and the overall low variation among our samples (biological coefficient of variation = 0.1748), indicates our FACS approach generated high quality cell-specific data.

### Disrupted meiotic X inactivation in both subfertile hybrids

We found disrupted meiotic X chromosome inactivation (diplotene cells) in both subfertile hybrids, but the disruption was more severe in ♀*mus*^CZII^ × ♂*dom*^WSB^ hybrids. Consistent with previous results (Larson *et al.* 2016, 2017), fertile *domesticus* and *musculus* males had very few X-linked genes expressed in diplotene cells. In contrast, both F1 hybrids had elevated expression of X-linked genes in diplotene cells (**Fig S2**), consistent with disrupted MSCI. In comparisons between F1 hybrids and their parents with the same X chromosome (♀*mus*^CZII^ × ♂*dom*^WSB^ vs. *musculus*; ♀*dom*^WSB^ × ♂*mus*^CZII^ vs. *domesticus*), F1 hybrids expressed more X-linked genes and every differentially expressed (DE) gene was overexpressed in hybrids. In contrast, there was no obvious asymmetry in expression on the autosomes (**Figs 1A, B**). When we compared the two hybrids, ♀*mus*^CZII^ × ♂*dom*^WSB^ had higher X-linked expression and DE genes between the hybrids were largely overexpressed in ♀*mus*^CZII^ × ♂*dom*^WSB^ hybrids (**Fig 1C**), suggesting that disrupted MSCI was more severe in ♀*mus*^CZII^ × ♂*dom*^WSB^ hybrids.

**Fig 1.**
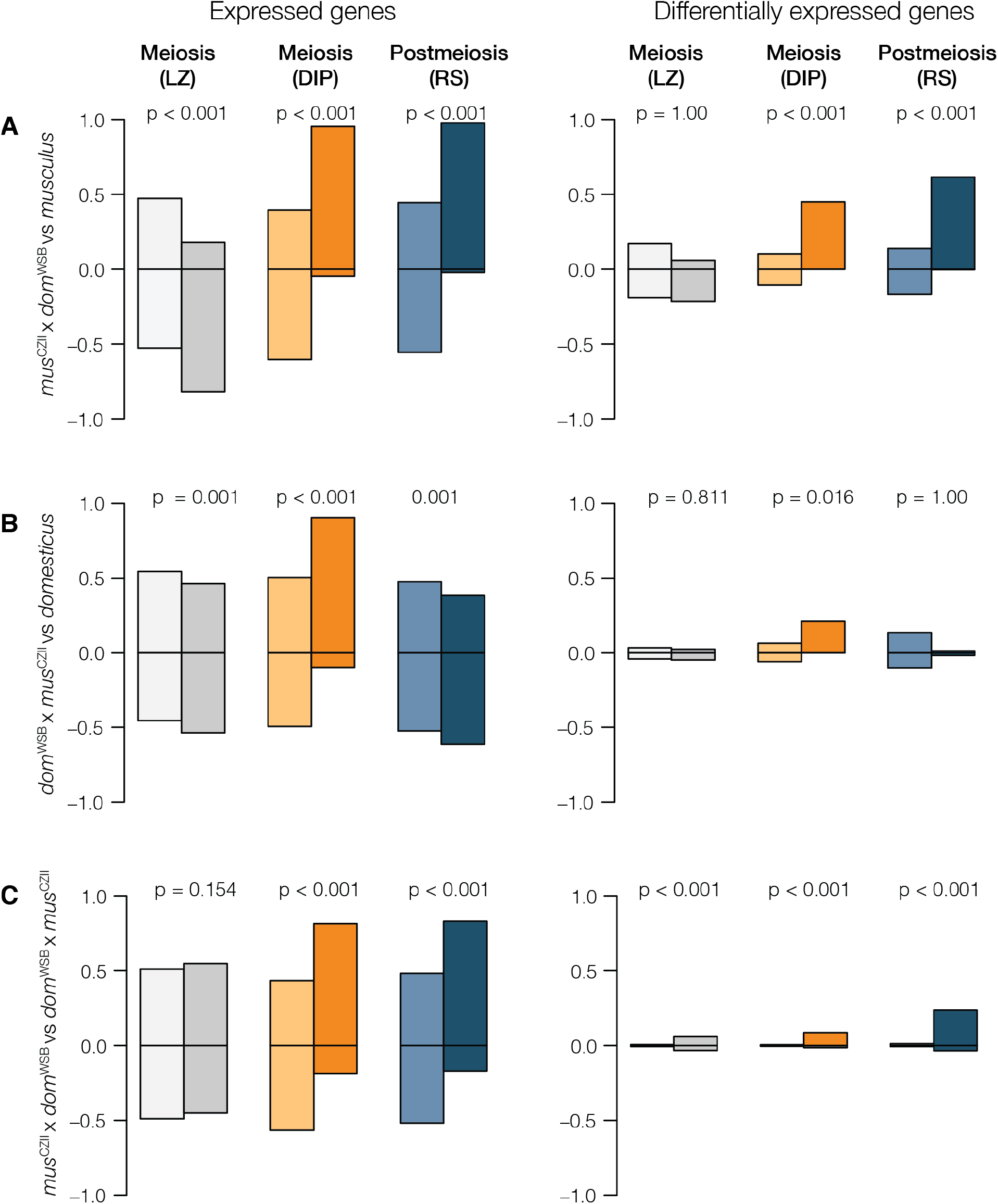
Gene expression comparisons among hybrids and parents for autosomes (light colors) and the X chromosome (dark colors). There are three contrasts: **A)** ♀*mus*^CZII^ × ♂*dom*^WSB^ hybrids compared to *musculus*, **B)** ♀*dom*^WSB^ × ♂*mus*^CZII^ hybrids compared to *domesticus*, **C)** ♀*mus*^CZII^ × ♂*dom*^WSB^ hybrids compared to ♀*dom*^WSB^ × ♂*mus*^CZII^ hybrids. The first column shows the proportion of genes with higher or lower expression in a given contrast out of the total genes expressed in each cell type. The second column shows the proportion of those genes that are DE. Significant p-values indicate contrasts where there was a significant difference in the proportion of over or underexpressed genes on the X chromosome compared to the autosomes (Pearson’s chi square test with FDR corrected p-values, Benjamini and Hochberg 1995). LZ = leptotene/zygotene cells (meiosis before MSCI), DIP = diplotene cells (meiosis after MSCI), RS = round spermatids (postmeiosis).

### Reciprocal hybrids have identical *Prdm9* genotypes

We characterized all four strains for allelic variation within Exon12 of *Prdm9* and confirmed that both *musculus* strains (PWK and CZECHII) shared the same *msc1* ‘sterile’ allele and both *domesticus* strains (WSB and LEWES) shared the same ‘sterile’ *dom3 Prdm9* allele (Forejt *et al.* 2021). Therefore, both ♀*mus*^CZII^ × ♂*dom*^WSB^ and ♀*dom*^WSB^ × ♂*mus*^CZII^ hybrids have the same *Prdm9* genotype at Exon12 (*msc1*/*dom3*).

### Imbalanced *Sly* and *Slx/Slxl1* copy numbers in reciprocal hybrids

We estimated gene copy number for postmeiotic amplicon families in our mouse strains using two methods and found that *musculus* had higher copy number for *Sly*, *Slx*, and *Slxl1* (**Table 2**). Our copy number estimates for *Sly* and *Slx* differed from what has been estimated using qPCR (Ellis *et al.* 2011) - we found higher copy numbers of *Sly* and lower copy numbers of *Slx*. Our estimates were closer to those from other studies that have used a computational approach to estimate copy number (Morgan and Pardo-Manuel de Villena 2017) and were similar to estimates for the *domesticus* Y chromosome assembly (Soh *et al.* 2014). Both our results and these other studies consistently found higher copy numbers in *musculus*, indicating there is an imbalance in *Sly* and *Slx/Slxl1* copy numbers of F1 hybrids relative to parental strains.

**Table 2:**
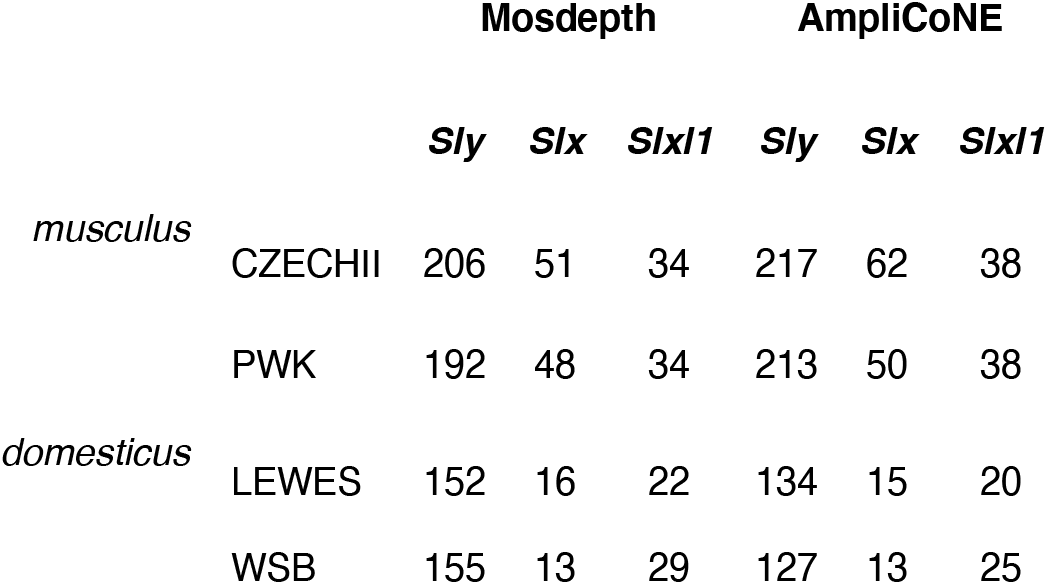
Copy number estimates for the *Sly* and *Slx/Slxl1* gene families for the wild-derived mouse strains used in this study.

### Postmeiotic disruption in *Sly*-deficient hybrids

The X chromosome was overexpressed in postmeiotic round spermatids of ♀*mus*^CZII^ × ♂*dom*^WSB^ hybrids (*Sly*-deficient), but not in ♀*dom*^WSB^ × ♂*mus*^CZII^ hybrids (*Slx*-deficient). Nearly all of the X-linked postmeiotic genes in ♀*mus*^CZII^ × ♂*dom*^WSB^ hybrids were overexpressed relative to *musculus* and more than half of these were DE (**Fig 1A**). In contrast, ♀*dom*^WSB^ × ♂*mus*^CZII^ hybrids had genes that were both over- and under-expressed relative to *domesticus* and DE genes tended to be expressed at lower levels (although this pattern wasn’t significant) (**Fig 1B**). There were no clear asymmetries on the autosomes for either hybrid relative to their parent. Given that both hybrids showed some degree of disrupted expression in meiotic cells, this suggests postmeiotic disruption is not a simple downstream consequence of earlier MSCI disruption, but is either an independent mechanism for postmeiotic disruption in *Sly*-deficient hybrids or there is a threshold of disrupted MSCI required to lead to downstream disruption.

SLX/SLXL1 and SLY compete for interaction with SSTY1 at promotors to regulate a suite of postmeiotic multicopy genes, including autosomal gene families *α-takusan* and *Speer* (Moretti *et al.* 2017, 2020). To test if we could detect misexpression of these autosomal gene families in our *Sly*-deficient hybrids, we used a sliding gene-window analysis (250 genes/window) to identify genomic regions with clusters of over or under-expressed genes between our reciprocal F1 hybrids. We found two small gene windows on chromosomes 5 and 8 that exceeded our threshold for overexpressed gene-windows (99th quantile modeled with a Poisson distribution) (**Fig S3**), but these regions did not overlap with any known multicopy gene families. We did not detect any large gene-windows that were consistently overexpressed in *Sly*-deficient hybrids as we did in crosses between *mus*^PWK^ x *dom*^LEW^ (Larson *et al.* 2017).

### Comparison of patterns of disrupted X expression across different hybrid genotypes

Finally, we used previously published data from Larson et al. 2017 to compare overlap in X-linked DE genes between reciprocal subfertile hybrids in this study (♀*mus*^CZII^ × ♂*dom*^WSB^, ♀*dom*^WSB^ × ♂*mus*^CZII^) with other subfertile hybrids (♀*mus*^PWK^ × ♂*dom*^LEW^). We found the greatest number of X-linked DE genes in postmeiotic round spermatids of ♀*mus*^CZII^ × ♂*dom*^WSB^ hybrids, and many of these same genes were also DE in ♀*mus*^PWK^ x ♂*dom*^LEW^ hybrids (**Fig 2**). The second highest number of X-linked DE genes were in meiotic cells (diplotene) of *mus*^PWK^ x *dom*^LEW^ hybrids, and a subset of these genes were also DE in ♀*mus*^CZII^ × ♂*dom*^WSB^ hybrids. There were approximately half as many DE meiotic genes in ♀*dom*^WSB^ × ♂*mus*^CZII^ hybrids, but nearly all of these were also misexpressed in the meiotic cells of the other two *mus* x *dom* hybrids. There were very few X-linked DE genes in the postmeiotic cells of ♀*dom*^WSB^ × ♂*mus*^CZII^ hybrids, though these genes did tend to overlap with DE postmeiotic genes in the other two subfertile hybrids.

**Fig 2.**
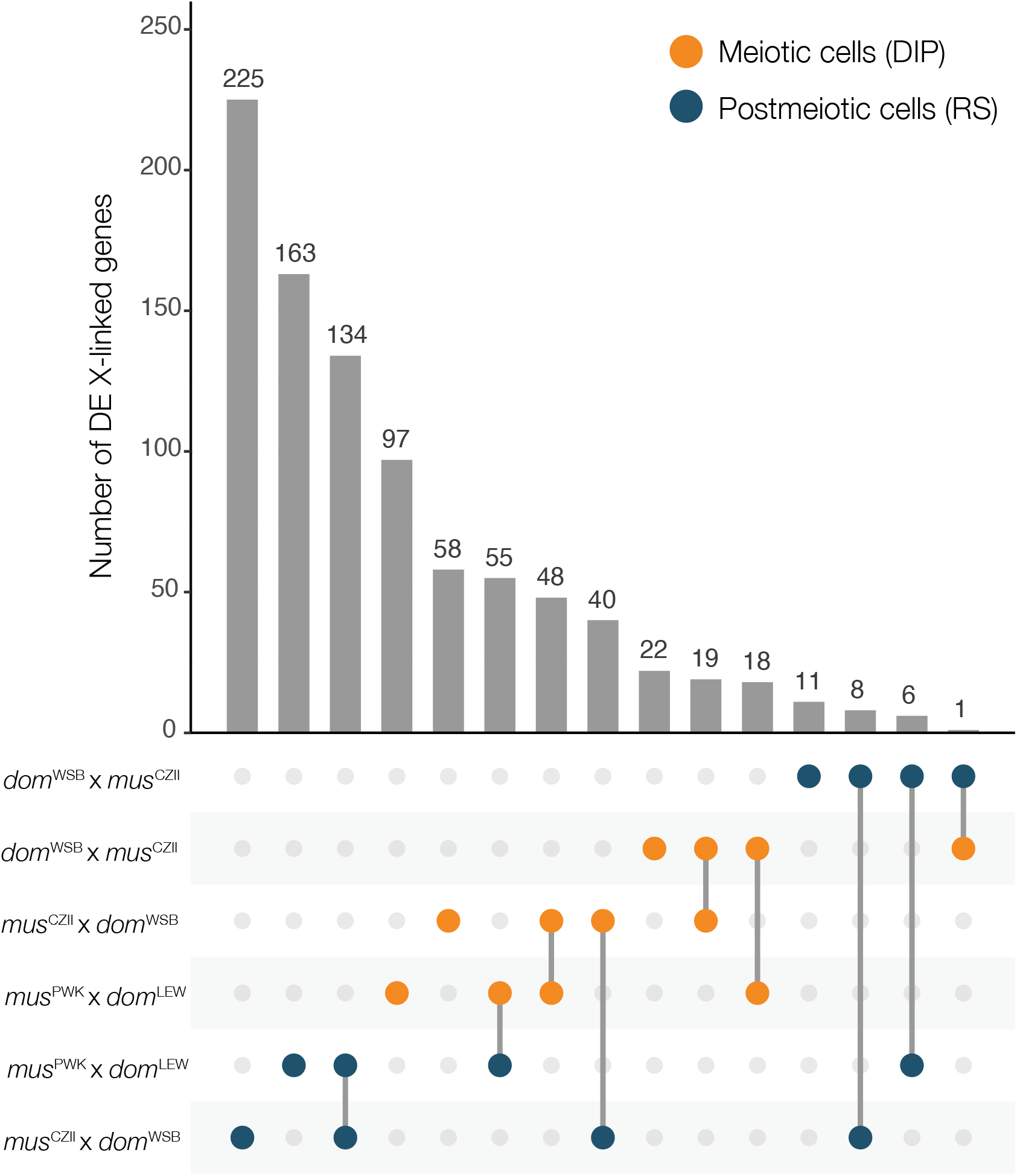
Number of X-linked DE genes across multiple subfertile hybrids. The dots indicate a contrast between a subfertile hybrid and its respective parental X chromosome and the barplot indicates the number of X-linked DE genes in that contrast. When there are two contrasts listed and a line connecting them it indicates the number of DE X-linked genes that are overlapping between the two contrasts. DIP = diplotene cells (orange), RS = round spermatids (blue).

## DISCUSSION

### Disrupted meiotic X inactivation in reciprocal F1 hybrids

Disruption of spermatogenesis during early meiosis has been linked to PRDM9, a protein that directs the location of meiotic recombination (Mihola *et al.* 2009). Divergence at PRDM9 DNA-binding sites can lead to incomplete meiotic synapsis of homologous chromosomes (Bhattacharyya *et al.* 2013; Davies *et al.* 2016; Gregorova *et al.* 2018), and associated disruption of MSCI (Good *et al.* 2010; Bhattacharyya *et al.* 2013; Campbell *et al.* 2013; Turner *et al.* 2014; Mack *et al.* 2016; Larson *et al.* 2017). We found disrupted MSCI in reciprocal subfertile hybrids, consistent with *Prdm9*-associated sterility in both F1 hybrids. Overall, the disruption was greater in ♀*mus*^CZII^ × ♂*dom*^WSB^ hybrids, but meiotic arrest was not complete in either cross, suggesting variation in the mechanisms that contribute to *Prdm9*-associated sterility.

PRDM9 defines where meiotic recombination will occur by adding histone marks that guide SPO11 protein to induce double-strand breaks, which are repaired as either crossovers or non-crossovers (Baudat *et al.* 2010; Myers *et al.* 2010; Parvanov *et al.* 2010). The C-terminal zinc finger domain of PRDM9 determines its binding affinity to a particular site, but *Prdm9* binding sites evolve very rapidly due to biased gene conversion. If one homolog has a mutation at a PRDM9 binding site, then PRDM9 will bind preferentially to the other homolog with the ancestral binding site, causing double strand break formation in only one chromosome. This break will be repaired using the mutated strand as a template, thus mutations at PRDM9 binding sites are rapidly incorporated into both homologs, leading to the erosion of PRDM9 binding sites over time (Myers *et al.* 2010; Baker *et al.* 2015). The same mechanism is what leads to autosomal asynapsis in hybrids (Smagulova *et al.* 2016; Davies *et al.* 2016; Gregorova *et al.* 2018). When hybrids are heterozygous at *Prdm9* and at PRDM9 binding-sites throughout the genome, PRDM9 binds preferentially to its ancestral binding sites, leading to asymmetric formation of double strand breaks, and the failure of autosomes to properly synapse. Asynapsed autosomes interfere with normal MSCI leading to the overexpression of the X chromosome, although the exact mechanism is still unknown (Forejt *et al.* 2021). Consistent with this model, we found reciprocal F1 hybrids both had disrupted MSCI and we found the same X-linked genes had disrupted meiotic X expression in both crosses, although there were slightly more disrupted genes in *mus*^PWK^ x *dom*^LEW^ hybrids (**Fig 2**). This suggests asymmetric PRDM9 binding occurs in both cross directions.

*Prdm9*-associated sterility is also influenced by an interaction with the *Hstx2* locus, a ~2.7 Mb region in the middle of the X chromosome (Storchová *et al.* 2004; Bhattacharyya *et al.* 2014; Lustyk *et al.* 2019). Complete meiotic arrest typically only occurs in F1 mice with a *musculus Hstx2* allele (*i.e., musculus* X chromosome), while F1 mice with a *domesticus* X chromosome may vary from subfertile to nearly fully fertile (Dzur-Gejdosova *et al.* 2012; Flachs *et al.* 2012; Mukaj *et al.* 2020). The *Hstx2* locus harbors a gene, *Meir1* that controls recombination rates, and is a strong candidate for directly modulating PRDM9 binding (Dumont and Payseur 2011; Balcova *et al.* 2016). This model predicts that sterility and disrupted gene expression will be the most severe in F1 hybrids with a *musculus* X chromosome. When we have examined expression in enriched cell populations, hybrids from *musculus* x *domesticus* crosses were subfertile and had disrupted MSCI (♀*mus*^CZII^ × ♂*dom*^WSB^, this study; *mus*^PWK^ x *dom*^LEW^, Larson *et al.* 2017), while some reciprocal hybrids were fertile with normal MSCI (*dom*^LEW^ x *mus*^PWK^, Larson *et al.* 2017). Indeed, we found that disrupted MSCI was much less severe in ♀*dom*^WSB^ x ♂*mus*^CZII^ hybrids (**Fig 2**), consistent with the idea that the *musculus* X chromosome is required for more severe meiotic disruption.

The severity of sterility in *musculus* x *domesticus* crosses appears to depend on allelic variation at *Prdm9* (Chromosome 17) and/or *Hstx2* (X chromosome). The PRDM9 C-terminal zinc finger domain is composed of repeats that are polymorphic within each subspecies (Buard *et al.* 2014; Kono *et al.* 2014; Vara *et al.* 2019) and ‘fertile’ and ‘sterile’ alleles have been described in both *musculus* and *domesticus* (Flachs *et al.* 2012; Mukaj *et al.* 2020). The strains we used in this study appear to have identical *Prdm9* alleles to those that have been described as ‘sterile’ in other studies (Mukaj *et al.* 2020; see also Forejt *et al.* 2021). Thus, despite both reciprocal hybrids having identical *Prdm9* genotypes (*msc1*/*dom3*), ♀*mus*^CZII^ × ♂*dom*^WSB^ produce some sperm with normal morphology (Table 1), suggesting that other loci must modulate *Prdm9*-associated sterility in this cross. In *musculus* PWK x *domesticus* B6 hybrids with two ‘sterile’ Prdm9 alleles, partial fertility appears to be associated with allelic variation on the X chromosome (Flachs *et al.* 2014), possibly at the *Hstx2* locus. Allelic variation on the X chromosome may also explain why complete meiotic arrest was not found in crosses with wild-derived strains in this study (♀*mus*^CZII^ × ♂*dom*^WSB^), by Larson et al. (2017; ♀*mus*^PWK^ x ♂*dom*^LEW^), and in some other *musculus* x *domesticus* crosses with two ‘sterile’ *Prdm9* alleles (Mukaj *et al.* 2020).

Allelic variation at *Prdm9* could also explain the subfertility of ♀*dom*^WSB^ × ♂*mus*^CZII^ hybrids, in the absence of the *musculus* X chromosome. F1 *domesticus* x *musculus* hybrids can be subfertile when both *Prdm9* alleles are ‘sterile’ (Flachs *et al.* 2012). The combination of two sterile *Prdm9* alleles and heterozygous PRDM9 binding sites throughout the genome may be sufficient to disrupt MSCI in ♀*dom*^WSB^ × ♂*mus*^CZII^ hybrids. However, it is unclear why MSCI would be disrupted in ♀*dom*^WSB^ × ♂*mus*^CZII^ hybrids, but not ♀*dom*^LEW^ × ♂*mus*^PWK^ hybrids, which also have two sterile *Prdm9* alleles. It is also unknown to what extent *Prdm9* contributes to the sterility phenotypes in ♀*dom*^WSB^ × ♂*mus*^CZII^ hybrids, given that other autosomal sterility factors have been mapped in ♀*dom*^WSB^ × ♂*mus*^CZII^ hybrids to chromosomes 2, 8 and 9 (Larson *et al.* 2018b).

In addition to allelic variation, the outcome of *Prdm9*-associated sterility is likely to be variable across cells within an individual. *Prdm9*-induced autosomal asynapsis is a threshold response. If a sufficiently large number of autosomes fail to pair (asynapsis rates > 60%) it leads to full meiotic arrest, while lower rates of asynapsis may lead to intermediate levels of meiotic disruption (Bhattacharyya *et al.* 2013; Mukaj *et al.* 2020). While MSCI is disrupted in ♀*mus*^CZII^ × ♂*dom*^WSB^ hybrids, the X chromosome still had lower expression in meiosis compared to other cell types (**Fig S2**) and a similar pattern was found for disrupted MSCI in ♀*mus*^PWK^ x ♂*dom*^LEW^ hybrids (Larson *et al.* 2017). This suggests cell-to-cell variation in the occurrence or magnitude of disrupted MSCI which may contribute to the range of sperm morphologies found in these hybrids - from severely impaired to apparently normal (**Table 1**).

### Asymmetric disruption of postmeiotic expression suggests genotype-specific hybrid sterility regulatory phenotypes

Postmeiotic disruption of X chromosome expression was observed in ♀*mus*^CZII^ × ♂*dom*^WSB^ hybrids but not in reciprocal♀*dom*^WSB^ × ♂*mus*^CZII^ hybrids (**Fig 1A, 1B**). Both F1 hybrids had earlier disruption of MSCI, which suggests that postmeiotic overexpression of the X chromosome is not a simple downstream consequence of disruption at earlier developmental timepoints. It is possible that downstream postmeiotic disruption depends on the magnitude of disrupted MSCI, or the disruption of particular X-linked genes or gene networks. Consistent with this, ♀*mus*^CZII^ × ♂*dom*^WSB^ hybrids had a higher proportion of disrupted X-linked genes in meiosis.

Asymmetric postmeiotic disruption is also consistent with antagonistic coevolution of X- and Y-linked multicopy gene families that leads to overexpression only in one cross direction. We found the X chromosome was overexpressed in postmeiotic cells of F1 hybrids that had a deficit of the Y-linked gene family *Sly* (♀*mus*^CZII^ × ♂*dom*^WSB^ hybrids), but not in reciprocal F1 hybrids that had a deficit of the X-linked gene family *Slx/Slxl1* (♀*dom*^WSB^ × ♂*mus*^CZII^ hybrids)(**Fig 1A, 1B**). *Sly* and *Slx/Slxl1* play a major role in suppressing or promoting postmeiotic expression of multi-copy sex-linked and interacting autosomal genes (Mueller *et al.* 2008, 2013, Kruger *et al.* 2019, Moretti *et al.* 2020). Imbalanced copy numbers of these genes in F1 hybrids may also disrupt postmeiotic expression networks. This could happen either independently of upstream meiotic disruption, or there may be an interaction among X-linked regulatory networks at different stages of development.

*Slx/Slxl1* originated from a single copy autosomal gene (*Sycp3*) that was transposed to the X chromosome (*Slxl1* then *Slx*) and eventually a copy emerged on the Y chromosome (*Sly*) (Kruger *et al.* 2019). Since their origin, these genes, additional sex-linked ampliconic genes, and associated autosomal ampliconic genes have undergone a massive co-amplification in different mouse lineages leading to divergent copy numbers in *domesticus* and *musculus* (see Table 2; Ellis *et al.* 2011; Turner *et al.* 2014; Soh *et al.* 2014; Morgan and Pardo-Manuel de Villena 2017). *Slx* and *Sly* appear to coevolve in a copy number arms race for interaction with SSTY1 at the promoter of thousands of postmeiotic genes (Moretti *et al.* 2020). Knockdown of *Sly* expression or duplications of *Slx/Slxl1* (i.e., *Sly*-deficient) leads to increased transmission of the X chromosome, abnormal sperm morphology, and upregulation of multicopy genes on the sex chromosomes (Cocquet *et al.* 2009, Kruger *et al.* 2019), as well as upregulation of the autosomal *Speer* (Chr 5) and *α-takusan* (Chr 14) gene families (Moretti *et al.* 2020). Knockdown of *Slx/Slxl1* expression (*i.e. Slx*-deficient) suppresses postmeiotic multicopy gene expression and leads to increased transmission of the Y chromosome and mild sperm abnormalities (Cocquet *et al.* 2010, 2012; Kruger *et al.* 2019). Reciprocal F1 hybrids between *domesticus* (*Sly* 130, *Slx/Slx1* 35, *Sly/Slx* ratio: 3.7) and *musculus* (*Sly* 215, *Slx/Slx1* 100, *Sly/Slx* ratio: 2.15) mirror these knockdown experiments: F1 hybrids from *musculus* × *domesticus* are *Sly*-deficient (130 *Sly,* 100 *Slx*, *Sly/Slx* ratio: 1.3) and F1 hybrids from *domesticus* × *musculus* are *Slx*-deficient (215 *Sly,* 35 *Slx*, *Sly/Slx* ratio: 6.1, **Table 2**).

Consistent with a *Sly/Slx* imbalance, we found *Sly*-deficient ♀*mus*^CZII^ × ♂*dom*^WSB^ hybrids overexpressed the X chromosome in postmeiotic cells (**Fig 1A**). We saw the same overexpression of the X chromosome in an independent contrast of *Sly*-deficient *mus*^PWK^ × *dom*^LEW^ hybrids (Larson *et al.* 2017). The same X-linked genes were overexpressed in both crosses, though there were more upregulated X-linked genes in ♀*mus*^CZII^ × ♂*dom*^WSB^ hybrids (**Fig 2**). In contrast, we found very few X-linked DE genes in *Slx*-deficient ♀*dom*^WSB^ × ♂*mus*^CZII^ hybrids, and there was no asymmetry in the expression of DE genes in ♀*dom*^WSB^ × ♂*mus*^CZII^ hybrids - genes were both up and down regulated relative to the *domesticus* X chromosome (**Fig 1B**). If anything, X-linked postmeiotic genes tended to be underexpressed in ♀*dom*^WSB^ × ♂*mus*^CZII^ hybrids relative to the *domesticus* X chromosome, but, unlike in *Slx*-deficient ♀*dom*^LEW^ × ♂*mus*^PWK^ hybrids (Larson *et al.* 2017), this pattern was not significant. Still, the handful of X-linked postmeiotic genes that were overexpressed in ♀*dom*^WSB^ × ♂*mus*^CZII^ hybrids were also upregulated in both *musculus* × *domesticus* hybrids (**Fig 2**).

In contrast to results from *mus*^PWK^ × *dom*^LEW^ hybrids (Larson *et al.* 2017), we did not find co-overexpression of ampliconic autosomal genes families (*Speer* or *α-takusan*) in ♀*mus*^CZII^ × ♂*dom*^WSB^ hybrids. The overexpression of these gene families in *Sly*-deficient hybrids was one of the strongest arguments for an independent mechanism of disrupted X expression in *mus*^PWK^ × *dom*^LEW^ hybrids. This lack of agreement makes it difficult to disentangle disrupted regulatory dynamics of *Sly* and *Slx/Slxl1* from possible downstream disruption of PRDM9 in this cross. However, the clear differences in postmeiotic expression between ♀*mus*^CZII^ × ♂*dom*^WSB^ hybrids and ♀*dom*^WSB^ × ♂*mus*^CZII^ hybrids indicates that postmeiotic disruption is genotype specific.

Whether or not postmeiotic sex chromosome overexpression contributes to sterility phenotypes in wild hybrids is still unknown. In knockdown studies, *Sly* and *Slx/Slxl1* have a major impact on sperm morphology (Cocquet *et al.* 2009, 2010, 2012; Kruger *et al.* 2019), but the extent to which these genes might contribute to hybrid sterility phenotypes in wild mice is still unclear (Campbell *et al.* 2013). In knockdown studies, *Sly*-deficient mice have severe sperm deformities (Cocquet *et al.* 2009) and biased X chromosome transmission, while *Slx*-deficient mice tend to have more typical sperm (but see Kruger *et al.* 2019) and biased Y chromosome transmission (Cocquet *et al.* 2010, 2012). The severe sperm deformities in *Sly*-deficient mice appear to particularly affect Y-bearing sperm, decreasing their mobility and providing a direct mechanism for how sperm morphology contributes to sex ratio skews (Rathje *et al.* 2019). However, it is still unclear if the imbalance manifested in mouse hybrids is sufficient to induce a regulatory misexpression phenotype. In F1 hybrids, the imbalanced copy number of *Sly* and *Slx/Slxl1* is certainly less severe in magnitude as total knockdown experiments. If there is a threshold of imbalance required for *Sly* or *Slx/Slxl1* to successfully outcompete the other (Moretti *et al.* 2020), we may not see the same impacts on sperm morphology or sex ratio distortion in wild hybrids. In general, we do find that *Sly*-deficient F1 *musculus* x *domesticus* hybrids have severely abnormal sperm morphology (see Table 1 and Larson *et al.* 2017), while *Slx*-deficient F1 *domesticus* x *musculus* hybrids tend to have more moderate sperm head abnormalities (Larson *et al.* 2017). Similar patterns have been observed in Y introgression lines that mismatch the *musculus* and *domesticus* X and Y chromosomes (Campbell and Nachman 2014). In this study, ♀*dom*^WSB^ × ♂*mus*^CZII^ hybrids also have severely abnormal sperm head morphology (Table 1), but there are clear autosomal contributions to these abnormalities (Larson *et al.* 2018b). To our knowledge, sex ratio distortion has not been documented in wild-derived crosses, though there is some evidence that it might occur in the mouse hybrid zone (Macholán *et al.* 2008, but see 2019).

## Conclusions

The elegant *Prdm9* incompatibility model is likely the single most important mechanism of F1 hybrid male sterility in house mice. We find evidence for *Prdm9*-associated disruption of meiosis in subfertile hybrids from reciprocal crosses of two wild-derived strains. We also find evidence that factors outside of *Prdm9* and *Hstx2* contribute to disrupted expression in F1 hybrids, providing support for the idea that hybrid sterility is a composite phenotype and likely polygenic (Campbell and Nachman 2014; Larson *et al.* 2017). Other factors such as autosomal incompatibilities and postmeiotic X-Y interactions are likely to be important contributions to overall hybrid sterility. Indeed, the variation we found in the extent and timing of disrupted X expression among different F1 hybrids may reflect interactions among disrupted meiotic and postmeiotic gene networks.

The mouse hybrid zone is a relatively recent contact that stretches across central Europe, with a fairly narrow width (Phifer-Rixey and Nachman 2015). Despite the recency of contact and the proximity of parental species, there are few F1 hybrids found in the center of the zone. Instead, the mouse hybrid zone is composed predominantly of advanced generation hybrids and backcrosses (Turner *et al.* 2012; Janoušek *et al.* 2012; Turner and Harr 2014), and hybrid males vary considerably in their fertility (Turner *et al.* 2012). *Prdm9*-associated sterility is strongest in an F1 background with a *musculus* X chromosome and depends on a combination of sterile *Prdm9* alleles (Forejt *et al.* 2021). Stretches of conspecific genomic regions, which are typical for backcrosses and advanced generation hybrids, can rescue meiotic synapsis (Gregorova *et al.* 2018). As a result, it is very unlikely that *Prdm9* alone can explain the reduced gene flow between *musculus* and *domesticus* in nature.

Studies of differential introgression in the mouse hybrid zone have consistently found the X chromosome to have restricted introgression, as well as a number of different autosomal regions (Tucker *et al.* 1992; Payseur *et al.* 2004; Macholán *et al.* 2007, 2011; Teeter *et al.* 2010; Janoušek *et al.* 2012; Turner and Harr 2014). Restricted introgression can point to regions of the genome that contribute to reproductive barriers. While there is some evidence for premating barriers between *musculus* and *domesticus* (Smadja and Ganem 2002, 2007; Bímová *et al.* 2011; Loire *et al.* 2017), the singular phenotype in all studies of these subspecies is the reduced fertility of hybrid males. Indeed, mapping studies have identified multiple regions of the X chromosome (Oka *et al.* 2004; Good *et al.* 2008a; Dufková *et al.* 2011; Turner *et al.* 2014; Turner and Harr 2014; Morgan *et al.* 2020) and numerous autosomal regions contributing to sterility in F1 hybrids (Larson *et al.* 2018b), F2 crosses and backcrosses (Good *et al.* 2008a; White *et al.* 2011; Turner *et al.* 2014; Schwahn *et al.* 2018; Morgan *et al.* 2020) and wild hybrids (Turner and Harr 2014). There is also evidence that XY mismatch contributing to abnormal sperm morphology (Campbell and Nachman 2014; Martincová *et al.* 2019b, 2019a), and patterns of directional introgression of the *musculus* Y chromosome into *domesticus* backgrounds (Macholán *et al.* 2008, 2019; Ďureje *et al.* 2012), consistent with postmeiotic X and Y chromosome conflict.

Complex hybrid incompatibilities, involving many genes, both autosomal and sex-linked, are a common feature of hybrid male sterility (Coughlan and Matute 2020). The multigenic nature of hybrid male sterility in house mice, and the availability of wild-derived strains makes this an excellent system to identify the genetic basis of hybrid sterility (Forejt *et al.* 2021) and relate these incompatibilities directly to reproductive isolation between natural populations.

## AUTHOR CONTRIBUTIONS

JMG conceived of the study and designed the experiments. ELL, EEKK, KEH, DV and SK performed experiments and collected data. ELL, EEKK, and KEH analyzed data. ELL and JMG wrote the manuscript with feedback from all co-authors.

## ACKNOWLEDGEMENTS

This work was funded by grants from the Eunice Kennedy Shriver National Institute of Child Health and Human Development of the National Institutes of Health (R01-HD073439, R01-HD094787 to JMG). ELL was supported by the National Science Foundation (DEB 1557059) and EEKK and KEH were both supported by the National Science Foundation Graduate Research Fellowship Program (EEKK: DGE-1313190, and KEH DGE-2034612). We also thank Pamela Shaw and the University of Montana Fluorescence Cytometry Core, supported by the National Institute of General Medicine Sciences of the National Institutes of Health (P30GM103338) for assistance with FACS and the University of Montana Genomics Core, supported by a grant from the M.J. Murdock Charitable Trust. Any opinions, findings, and conclusions or recommendations expressed in this material are those of the author(s) and do not necessarily reflect the views of the National Science Foundation or the National Institutes of Health.

## SUPPLEMENTARY DATA

**Table S1: Table of individual male reproductive phenotypes (.csv).** Table includes each individual mouse ID (*e.g.* CCPP 21.1M stands for dam x sire, litter number, individual number and sex; CC = CZECHII, PP = PWK, WW = WSB, LL = LEWES), cross type, dates the mice were born and phenotype, their age at phenotyping, measures of body size (weight, body length, tail length, right hind foot, left ear length), weights of paired testes and seminiferous vesicles, counts of motile and nonmotile sperm, counts of total sperm, and counts of sperm head morphology categories.

**Figure S1.**
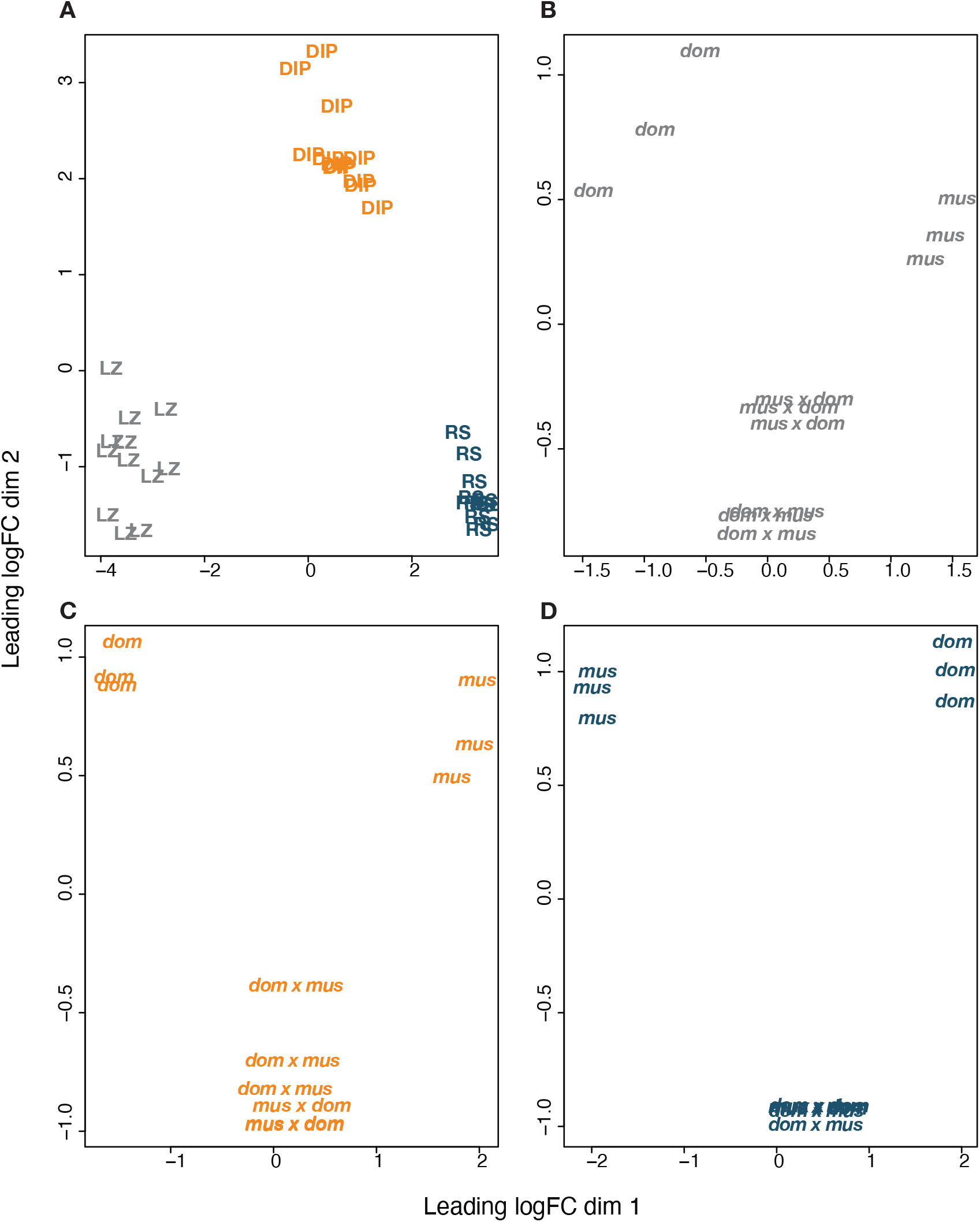
Clustering of gene expression profiles. Multidimensional scaling plots (MDS) of the Euclidean distance among gene expression profiles. Distance approximates the typical log2 fold changes between samples. **A)** RNAseq profiles cluster overall by cell type. LZ = leptotene/zygotene cells (gray), DIP = diplotene cells (orange), RS = round spermatids (blue). **B-D)** Within each cell type, RNAseq profiles cluster by subspecies, with F1 hybrids intermediate to the two parental subspecies. **B)** LZ = leptotene/zygotene cells (gray). **C)** DIP = diplotene cells (orange). **D)** RS = round spermatids (blue)

**FigS2.**
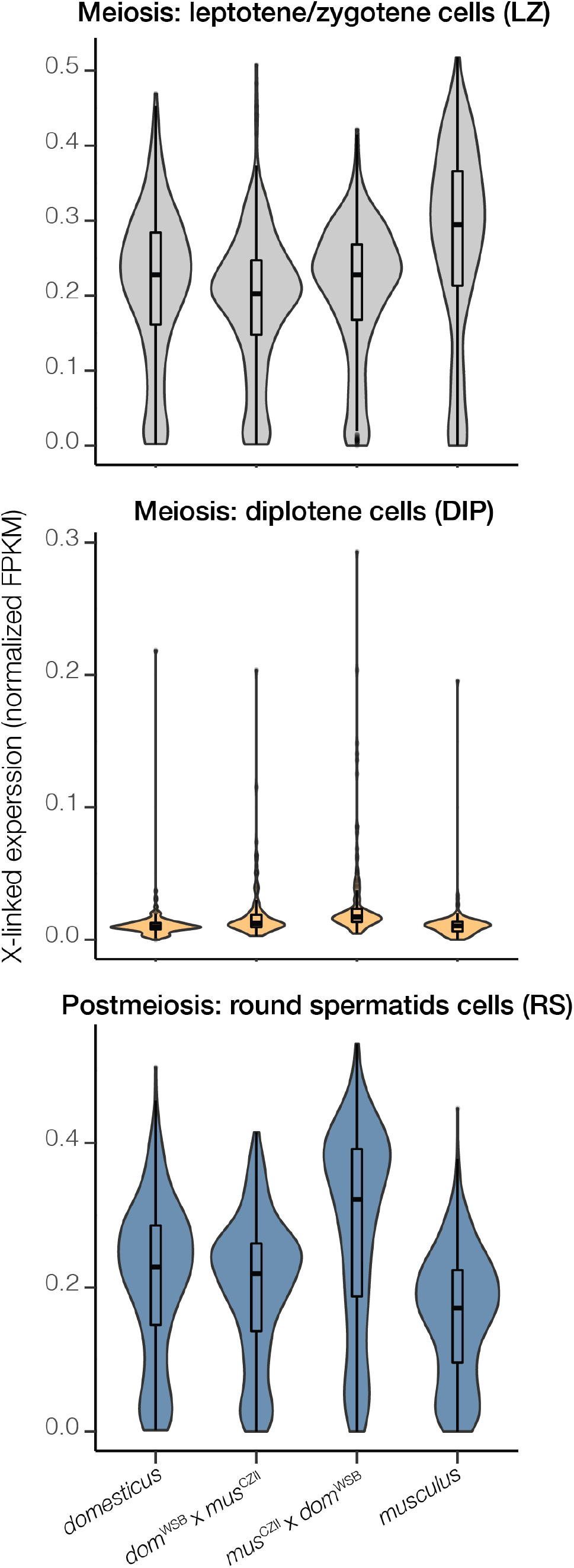
X chromosome expression across different cell types and crosses. The distribution of X-linked gene expression in normalized FPKM values (values range 0 to 1). The violin plots show the density of genes with a given expression level and the boxplots depict the median values and quartiles. Gene expression was restricted to genes that have an FPKM > 1 in at least 3 samples per cell type. X-linked expression was elevated in diplotene cells of both hybrids and in round spermatids of ♀*mus*^CZII^ × ♂*dom*^WSB^ hybrids.

**Figure S3.**
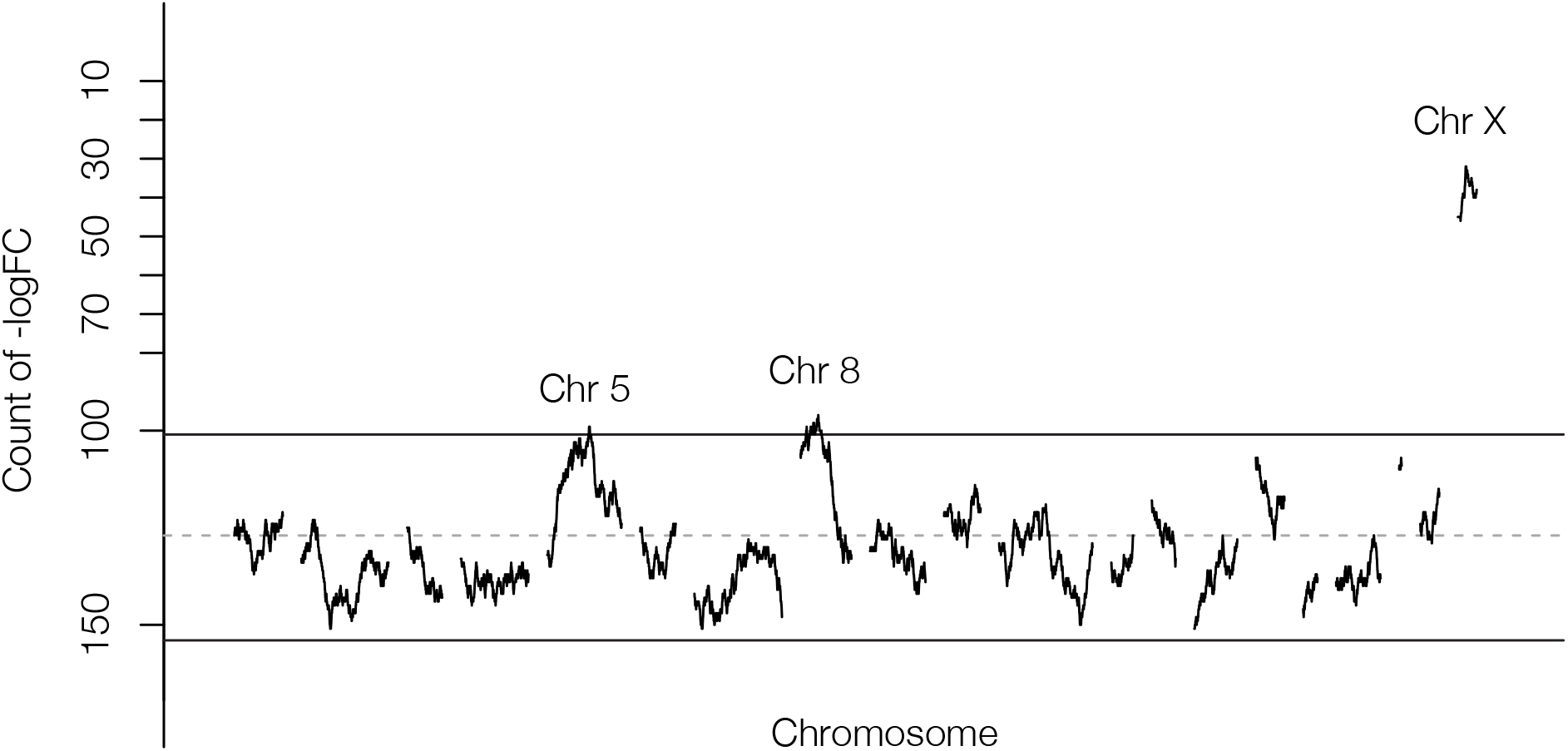
Spatial patterns of postmeiotic expression between subfertile hybrids. Sliding-gene windows (250 genes) for counts of underexpressed genes in postmeiotic cells (round spermatids) between ♀*mus*^CZII^ × ♂*dom*^WSB^ hybrids and ♀*dom*^WSB^ x ♂*mus*^CZII^ hybrids. Solid lines represent the 99th quantile modeled with a Poisson distribution. Note the Y-axis is plotted so that underexpressed genes fall below the 99th quantile and overexpressed genes are above the 99th quantile. Chromosomes 5 and 8 had relatively small windows of genes overexpressed in *Sly*-deficient ♀*mus*^CZII^ × ♂*dom*^WSB^ hybrids, but these windows did not coincide with known multicopy gene families (*Speer*/*α-takusan*).

## Notes

### Competing Interest Statement

The authors have declared no competing interest.

